# Robust network inference using response logic

**DOI:** 10.1101/547216

**Authors:** Torsten Gross, Matthew Wongchenko, Yibing Yan, Nils Blüthgen

## Abstract

**Motivation:** A major challenge in molecular and cellular biology is to map out the regulatory networks of cells. As regulatory interactions can typically not be directly observed experimentally, various computational methods have been proposed to disentangling direct and indirect effects. Most of these rely on assumptions that are rarely met or cannot be adapted to a given context.

**Results:** We present a network inference method that is based on a simple response logic with minimal presumptions. It requires that we can experimentally observe whether or not some of the system’s components respond to perturbations of some other components, and then identifies the directed networks that most accurately account for the observed propagation of the signal. To cope with the intractable number of possible networks, we developed a logic programming approach that can infer networks of hundreds of nodes, while being robust to noisy, heterogeneous or missing data. This allows to directly integrate prior network knowledge and additional constraints such as sparsity. We systematically benchmark our method on KEGG pathways, and show that it outperforms existing approaches in DREAM3 and DREAM4-challenges. Applied to a perturbation data set on PI3K and MAPK pathways in isogenic models of a colon cancer cell line, it generates plausible network hypotheses that explain distinct sensitivities towards EGFR inhibitors by different PI3K mutants.

**Availability and Implementation:** A Python/Answer Set Programming implementation can be accessed at github.com/GrossTor/response-logic.

## 1 Introduction

Complex molecular networks control virtually all aspects of cellular physiology as they transduce signals and regulate the expression and activity of genes. Understanding those molecular networks requires an appropriate simplification of the stupefying complexity that we find in cells. A very successful and common abstraction in molecular cell biology is to define effective modules and map out their interactions [22]. But even though new experimental techniques can reveal and quantify countless cellular components in ever increasing level of detail, they typically cannot identify the relationships between them. This is why for more than two decades various methods were developed to infer gene regulatory networks, signalling pathways and genotype-phenotype maps [12]. These methods vary widely in their notion of network (e.g. directed vs. undirected, weighted vs. unweighted links), their mathematical methodology (e.g. statistical measures vs. model-based parameter fits), or their goals (e.g. interaction discovery vs. network property characterization vs. perturbation response prediction) [33, 26, 11, 18, 25, 4, 34]. Not surprisingly, different methods produce radically different results on same data sets [32, 31]. This makes for an intricate choice of method and guarantees a certain degree of arbitrariness in interpreting the inferred networks.

One major goal of network inference for signalling and regulatory networks is to derive directed networks, that is, to infer information about causal relations within the studied system. This differs profoundly from the inference of undirected associations between node pairs, such as by correlation, as it requires to trace the flow of information through the network. A popular approach is to use time-series data, for which methods like convergent cross mapping [40, 8] or Granger causality [19] can distinguish correlation from causation, given sufficiently dense samples. But most often, experimental protocols or excessive expenditures preclude the observation of suitable temporal trajectories for many contexts in molecular biology. Thus, a complementary approach is to observe the system’s responses, for instance the steady state response, to a set of localized perturbations [7, 37, 43]. Depending on the specific system, these perturbations could, for example, be gene knockouts or kinase inhibitions. However, existing methods rely on context-specific assumptions whose validity is hard or impossible to assess in practice, which makes it very difficult to interpret their results. Facing this challenge, we asked whether we could derive a more generally applicable scheme for the inference of directed networks – a method that is based on a principle which is accurate enough for most contexts while also sufficiently simple to allow for an intuitive understanding of how the network structure was resolved. Furthermore, we noticed that even though most network inference problems are embedded within very well studied contexts, the vast majority of reverse-engineering methods predicts networks *de novo*. Therefore, we additionally aimed for a method that could readily incorporate prior knowledge about presence or absence of certain links or about other known network properties. This resulted in what we call the response logic approach.

In the following, we describe the response logic approach in more detail and then benchmark it by (i) assessing the performance using synthetic data derived from KEGG pathways [24], and (ii) comparing its performance to competing methods using the communitywide inference challenges (DREAM) [38]. Finally, we use the approach to study RAS/MAPK/PI3K signalling in a colon cancer cell line, and predict differences in the signalling network topology due to different PI3K mutants, that manifests in differential sensitivity of a colon cancer cell to various targeted drugs.

## 2 Method

We developed a method to infer directed network structures from perturbation data that we term response logic (see Figure 1).

**Figure 1:**
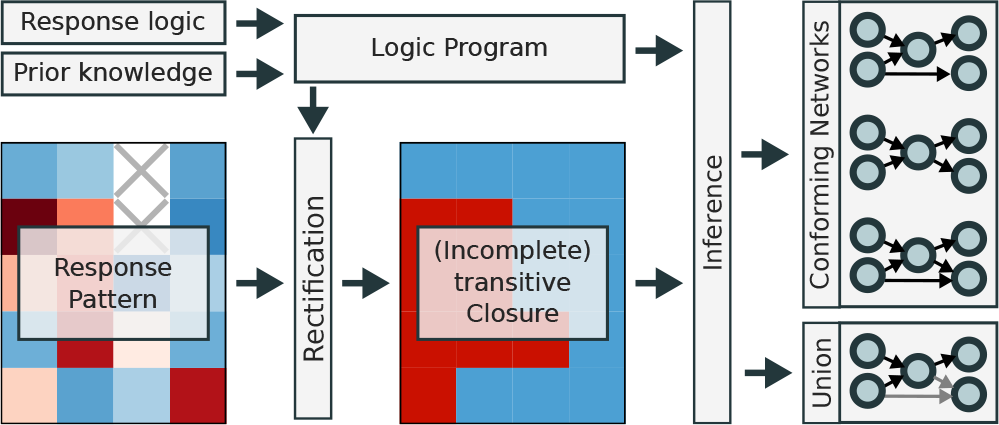
The steps of the response logic approach. The response logic and all additional prior network knowledge are formulated as a logic program. It is first used to rectify the experimentally determined response pattern, and second, takes the resulting (potentially incomplete) transitive closure as input to infer either all individual conforming networks or the union thereof.

As an input, this method requires binary information about which nodes in the network respond to which perturbation, together with a rank of confidence of each data point. We refer to this set of experimental observations as the response pattern. Given this information, the response logic approach infers networks that agree to the following simple rule: A perturbation at a node is propagated along all outgoing edges to the set of connected nodes, and these responding nodes will in turn propagate the signal and so forth. Consequently, a perturbation of a node causes a response at all nodes to which it is connected by a path, and no response at all others. The information about which node can be reached from which other nodes is known as the network’s transitive closure. Thus, the central assumption of our response logic approach is that experimentally observed responses are in agreement with the transitive closure. This assumption then leads to the inverse problem of identifying the networks whose transitive closure actually matches the response pattern.

The algorithm to infer these networks consists of two main steps. Using a logic programming approach, it first modifies the experimental response pattern to match a transitive closure (rectification step) and then infers either all individual networks that comply to the given data or the union over all those conforming networks. We will describe the different steps in the following sections.

### 2.1 Rectifying the response pattern

The response logic approach interprets the measured response pattern as a noisy, incomplete transitive closure. But because of misclassification, a response pattern might not match any actual (incomplete) transitive closure. Consider for example a three node network in which all nodes are observed to respond to a perturbation at node one. This implies two paths, from nodes one to two and nodes one to three. Therefore, if a perturbation at node two causes a response at node one, node three is expected to respond as well. But assume that this response at node three was not observed (misclassification). Then, there is no directed network with a transitive closure that would match this response pattern. We expect that such misclassification occurs rather often when working with experimental data because of experimental noise or because the system under consideration does not fully comply with the assumptions of the response logic. Thus, it is necessary to identify the most relevant subset of the response pattern that forms an (incomplete) transitive closure which can then be used to infer networks.

Our rectification algorithm requires to rank the observations of the response pattern from most to least confident. Typically, such confidence levels are readily available since the response pattern is often derived from a binarization of continuous experimental readouts, in which case a confidence score could be the distance to the binarization threshold, or a score of statistical significance. The algorithm then iterates the elements of the response pattern from high to low confidence, and at each step, determines whether the so far collected elements form a transitive closure and also conform with additional constraints from prior knowledge. This is done using a logic program (see below), which determines if there is any network that is compatible with these elements of a transitive closure. If the new element is compatible, it is added to the collection of conforming data and otherwise discarded. The more data points enter the collection the more restrictions apply to the remaining elements of the response pattern. As high confidence observations are taken into account first they are thus less likely to be discarded, ensuring that we extract the most relevant subset of the response pattern that indeed forms an incomplete transitive closure that is in line with additional heuristic constraints.

Figure 2 demonstrates this scheme for a toy network of five nodes, of which we assume that four nodes were perturbed (indicated as flashes in Figure 2A, top). The resulting response pattern then consists of a five by four matrix, and we assume that two data points are missing, and two elements of the response pattern do not match the transitive closure (compare Figure 2A middle and bottom). In Figure 2B, we exemplify how the response pattern is iteratively rectified. We assume that we know that a link from node two to node three exists and that there is no link between node one to node three (green stars). Given this prior knowledge, already the first (highest confidence) data point (yellow star in the leftmost panel), additionally implies that node three also responds to a perturbation of node one. Any subsequent data point that would be in conflict with this information will be discarded. The middle panels of Figure 2B show that the five most trusted data points constrain five other elements of the rectified response pattern. Amongst them are the two misclassified, as well as the two missing data points. Therefore, in this toy example, the high confidence data points automatically correct these false or missing pieces of information. The bottom panel of Figure 2B shows how adding data points increasingly constrains the network structure. Once all data is considered, most of the links (but not all, as discussed later on) are known to be either present of absent from the network. Note, however, that the rectification process does not require to compute the shown union of conforming networks, but only requires to determine if for any network at all, all constraints are satisfiable, which is computationally far less expensive.

**Figure 2:**
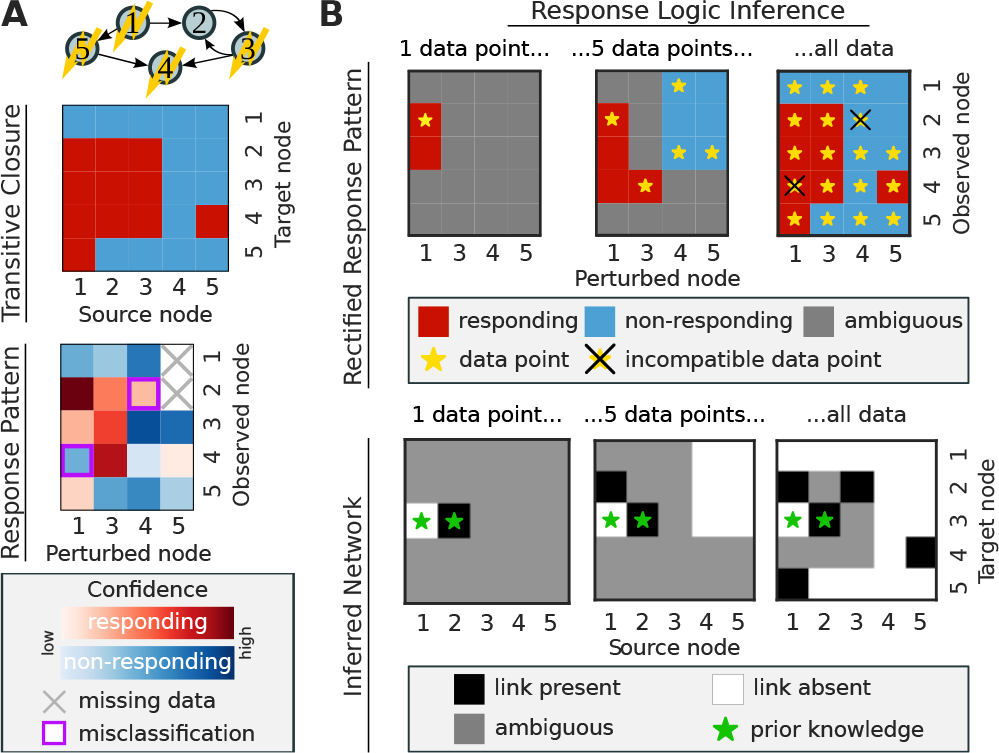
Response logic inference of a toy network. **A** From top to bottom: An example network of five nodes,where flashes indicate which nodes were perturbed; the full transitive closure; the response pattern that captures parts of the the network’s transitive closure, with missing or misclassified data and confidence scores. **B** Three steps during data rectification: Data points are added sequentially from high to low confidence (stars in top row), and increasingly constrain the inferred network and the (rectified) response pattern (red and blue fields in top row). Bottom row shows the inferred network at that step during rectification.

### 2.2 Finding conforming networks with logic programming

Mapping the response pattern to its corresponding set of conforming networks is a substantial computational challenge, as there are 2^*N ·N*^ possible directed networks (with *N* being the number of nodes), making it infeasible to enumerate all networks even for small sizes. We therefore solve the search problem with a logic programming approach, which is a form of declarative programming where the problem is represented via a set of logical rules. We chose to use the logic modelling language Answer Set Programming (ASP) [3], as implemented in the Potsdam Answer Set Solving Collection [17]. For ASP solving, we apply the *clingo* [16] system.

ASPs generate-define-test pattern [30] allows for a convenient encoding of the response logic, and the specific encoding is detailed in Supplementary Material S1. In short, we generate the collection of answer sets, consisting of all possible network structures, then define auxiliary predicates, in our case the networks transitive closure, and then test whether this transitive closure agrees with the data and also whether the tested network complies to all other heuristic constraints. Then the ASP solver, *clingo*, allows to enumerate all conforming networks. Note that computational effort needed to identify a conforming network heavily depends on network size and the provided heuristic constraints. But overall, the logic programming approach infers networks of up to 100 nodes within seconds, without any parallelization.

The previously discussed data rectification sequentially checks the satisfiability of every data point and could therefore become a performance bottleneck for large systems. However, because this process only requires to decide whether any network at all is in agreement with the latest data, instead of having to provide the entire set of conforming networks, we can solve a much simpler logic program. Its crucial performance gain arises from avoiding to generate an answer set for each possible network structure. The specific implementation is described in Supplementary Material S1.

### 2.3 Identifiability and heuristic constraints

While every directed network has a single transitive closure, a transitive closure can often be mapped to many different networks, even more so if the transitive closure is only partially known. Thus, we can usually not infer a unique directed network from a rectified response pattern alone. For example, any feedback loop creates a strongly connected network component, that is, a set of nodes for which any pair is connected by a path. Therefore their response pattern is independent on how exactly they are connected to each other. Similarly, the response pattern does not change with any additional feed-forward loops that cuts short an existing path. To resolve such structures we need to resort to additional constraints that are derived from contextual knowledge about the studied system. A crucial advantage of the response logic approach is that it can easily integrate various kinds of such constraints. Here, we want to exemplify this and introduce those constraints that are used in the applications shown further below.

Rarely will we analyse networks that have never been studied before and therefore one can use prior knowledge to constrain networks, such as by requiring the presence of well-established links in the network, or by excluding links that are physically not feasible (such as interactions between molecules that are not in the same compartment). This information can directly be integrated into the logic program by defining the presence or absence of links as additional constraints. In addition, the logic program can also accommodate more subtle constraints, such as to enforce bounds on the numbers of incoming and outgoing edges of (groups of) nodes, see the implementation in Supplementary Material S1. This allows, for example, to encode the information that a module of nodes signals to other parts of the network without having to explicitly state which of the module’s nodes has the outward link. The same idea holds for a module that is known to receive at least one input to any subset of its components. Note that these types of constraints directly limit the space of possible networks and in turn that of the transitive closures. They will thus influence how the response pattern can be rectified and must be taken into consideration during the process.

But even these additional constraint might not sufficiently limit the number of conforming networks to consider them individually. Alternatively, an extension of the logic program, described in Supplementary Material S1, allows to efficiently find the union of all answer sets. This union reveals which links (or missing links) appear in all solutions and those links where there is ambiguity. Exposing the ambiguous parts of the network is particularly valuable because it can either guide the choice of additional perturbation experiments or reveal effective strategies on how to further filter the set of solutions.

One widely used strategy in this regard is to require an overall sparse architecture [44]. We would thus want to identify the conforming networks with the fewest links. However, naively parsing all network solutions will be infeasible when the set of solutions is large. To overcome this problem we developed an algorithm that sequentially removes as many ambiguous links as possible, without violating any constraint. To do so, after every link removal the pruned network is tested for satisfiability. If it complies to the given constraints, the link remains removed and the procedure continues. Otherwise the link is considered necessary and the procedure continues without the removal of the link. This leads to what will be referred to as the *sparsified network*. Yet, such scheme is only reasonable if the order by which links are removed, reflects to some extent a knowledge about which links are more likely to be absent in the underlying network, and should therefore be tested for removal first. However if such information is not available, one can use yet another approach to filter for sparse networks, termed the *parsimony constraint*. This constraint asks whether a link from a conforming network can be removed without it changing the network’s transitive closure. If that is the case, the network is considered non-sparse and is removed from the solution set. The specific encoding is found in Supplementary Material S1. While this procedure does not generally single out a unique solution as before (multiple networks can be parsimonious), it was nevertheless observed to drastically reduce the solution space.

Taken together, a response pattern will typically be compatible with a large number of network topologies, but various types of prior network information can be incorporated into the response logic approach to reveal a finer network structure than what would have been possible from the response pattern alone. At the same time, the approach states explicitly whether or not the presence or absence of a link can be inferred from the given data and constraints.

### 2.4 Implementation and data acquisition

The response logic approach is implemented in Python 3.6 as a package available at github.com/GrossTor/response-logic. Numerical computations, data handling and plotting was done using the `SciPy` libraries [23] and `seaborn`. Additional functions were taken from the networkx package [21]. Clingo’s python API (version 5.2.2) [16] is used to ground and solve the Answer Set Programs.

The repository contains all Answer Set Programs, which are accessed by the the main *response.py* module. It includes the *prepare ASP program*-function to set-up a logic program according to provided data and additional constraints, the *conform response pattern*function that rectifies the response pattern, as well as various functions to solve a logic program. Additionally, a repository available at github.com/GrossTor/response-logic-projects includes all scripts and data that were used to obtain the results from the following sections.

The provided KEGG data [24] was retrieved via the `KEGG` package within the `biopython` library. The KEGG pathway maps database was parsed for human pathways and the retrieved KGML-files were used to build network representations based on their “relation elements”.

The data and evaluation scripts for the DREAM3 and DREAM4 challenge was retrieved with the official `DREAMTools` python package [9]. Leader-boards were taken from www.synapse.org/#!Synapse:syn3049712/wiki/74631 and Figure 3 in [31].

**Figure 3:**
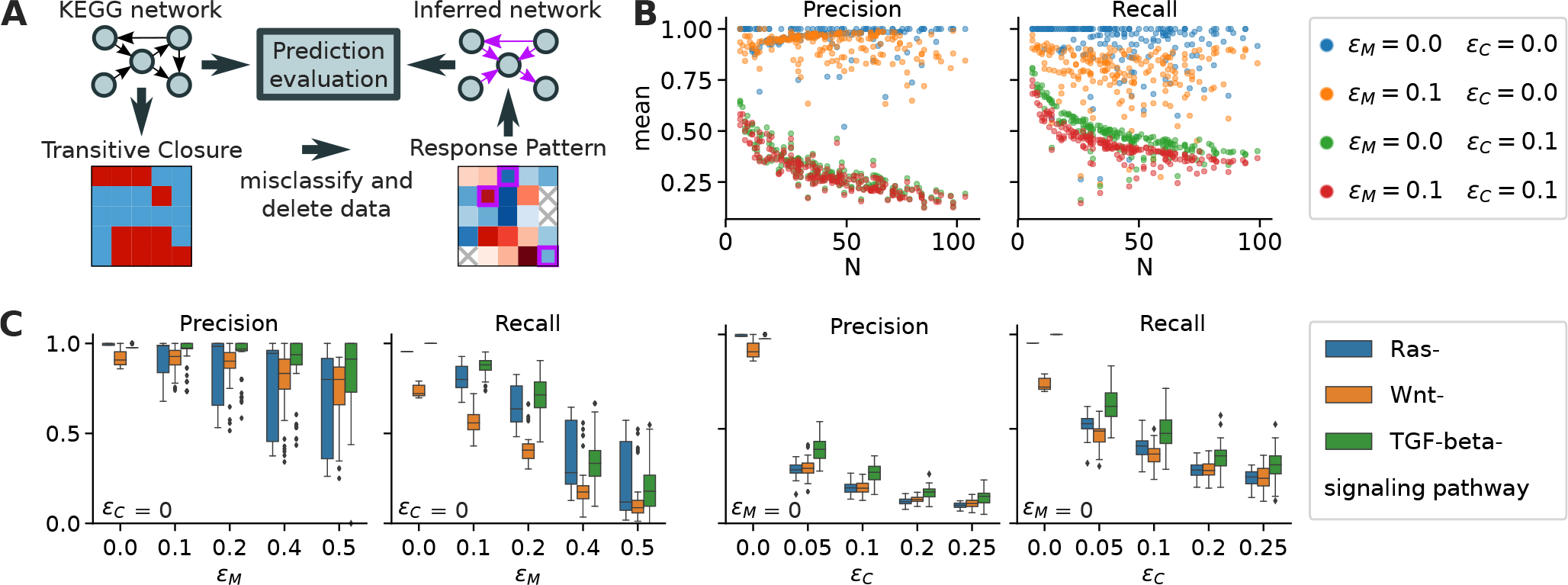
The performance of the response logic approach on synthetic data generated from 270 human KEGG pathways [24]. *N* denotes network size, *ϵ_M_* quantifies the fraction of missing and *ϵ_C_* the fraction of misclassified data points. **A** Data generation and scoring scheme. **B** Each dot per color represents a different pathway, colors represent different parameters for misclassification (*ϵ_C_*) and missing data (*ϵ_M_*). **C** Precision and recall for three particular signalling pathways as a function of the fraction of misclassified or missing data.

The SW-48 perturbation data was generated using a SW-48 cell line, and two derived clones with mutations in PI3K. Cell lines were obtained from Horizon Discovery. All lines were maintained in RPMI (Invitrogen) with 10% FBS (Invitrogen). Cell growth was assessed using the Cell Titer 96 Aqueous One Solution Cell Proliferation Assay (Promega). Cells were treated with compound 24 hours after plating and grown for 72 hours. The cell growth was determined by correcting for the cell count at time zero (time of treatment) and plotting data as percent growth relative to vehicle (DMSO) treated cells. Reverse Phase Protein Array (RPPA): Cells were treated 24 hours after plating and incubated with inhibitor (GDC0973, GDC0068, Erlotinib) or solvent control (DMSO) for 1 hour, and then stimulated either with EGF, HGF, and IGF or with control (BSA) for 30 min. Cells were lysed in T-PER (Thermo), 300mM NaCl, cOmplete^®^ protease in-hibitor (Roche), and Phosphotase Inhibitor Cocktails 2,3. RPPA measurements were carried out by Theranostics Health.

## 3 Results

### 3.1 Performance assessment on KEGG pathways

We first set out to systematically quantify how misclassification and missing data in the experimentally determined response pattern impacts the quality of the predicted network structure. To this end, we inferred network structures from synthetic data sets. As a relevant and representative collection of test networks, we extracted all 270 human gene regulation and signal transduction networks (maximally containing 100 nodes) from the KEGG pathway database [24]. For each of these network structures we generated its transitive closure, which we considered as the immaculate response pattern. Then, we repeatedly generated a random confidence pattern, *C*, where each entry is drawn from a uniform distribution between zero and one. To evaluate the effect of missing data, we remove a fraction *ϵ_M_* of data points from the perfect response pattern and to evaluate the effect of measurement error, we also misclassify a fraction *ϵ_C_* of the remaining data points. Missing or misclassified data points were chosen with a probability that was proportional to their confidence score *C_ij_*. We then used the resulting response and confidence patterns to infer the sparsified network, as defined in the previous section, via the response logic approach and, comparing it to the original KEGG network, computed precision and recall as performance scores, see Figure 3A.

For each of the 270 KEGG networks the procedure was repeated 50 times for different choices of *ϵ_M_* and *ϵ_C_*, and the mean of the scores is shown in Figure 3B. In the absence of misclassifications (*ϵ_C_* = 0, blue and orange dots in Figure 3B), prediction errors stem exclusively from the previously discussed multitude of conforming network structures. Interestingly, for a vast set of biological pathways the resulting inference errors are rather mild, and highly accurate predictions can be made independent of network size. However, once misclassifications are present, the predictivity is markedly reduced. Interestingly, this effect increases with growing network size.

We next examined the dependency on missing data and misclassification rates in more detail for the three signalling pathways RAS, Wnt and TGF-beta. We chose to scan the parameters from 0.0 to 0.5 and 0.0 to 0.25 for *ϵ_M_* and *ϵ_C_*, respectively, as a complete loss of information would either occur when all data was missing, *ϵ_M_* = 1, or half of the entries were misclassified, *ϵ_C_* = 0.5 (*ϵ_C_* = 1 would produce an inversion of the response pattern). For all pathways, we found that recall is more affected by missing data than precision (see Figure 3C). That is, with less data the predicted links remain rather accurate but fewer of them are predicted. We also confirmed our previous finding that misclassification reduces prediction scores much stronger than missing data. Interestingly, even when half the data was discarded, in many instances precision remained still close to one. This suggests that discarding lowconfidence data points rather than risking to accept many misclassified data points might be a good strategy to improve predictions. We will re-examine this idea by the end of the next section.

### 3.2 Response logic approach outperforms competing methods in DREAM challenges

The Dialogue for Reverse Engineering Assessments and Methods (DREAM) [38] provides community-wide reverse-engineering challenges that foster the development of new systems biology models. Particularly, the DREAM3 and DREAM4 in-silico challenges [31, 20] assessed the performance of various gene networkinference methods to which we can compare the response logic approach. In these two challenges various biologically plausible *in silico* gene networks of different sizes were simulated under stochastic conditions to emulate realistic transcription dynamics resulting from knockdowns and knockouts of each single gene. Participants were given the resulting time courses, the steady states, and the wild type level of each gene and asked to infer the directed network structure from them. A ranked list of predicted gene pair interactions was then compared against the gold standard from which the area under the precision-recall and the area under the Receiver Operating Characteristic curve are computed, see Supplementary Material Figures S1 and S2. Comparing these to a null model provides p-values for each of the given five networks per network size that then get combined into a single overall score [39].

To infer the DREAM networks with the response logic approach we generated response patterns from the *in silico* knockout experiments of these challenges only (not considering knockdown or time series data). These were computed as follows. When *K_ij_* denotes the level of gene *i* after knockout of gene *j*, and the wild type levels are **w**, we defined the normalised global response matrix, *R*, as

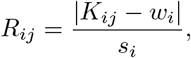

with *s_i_* being the standard deviation of the knockout levels of gene *i* (row *i* of *K*). We then defined gene *i* to be responding to a knockout of gene *j* if *R_ij_* > 1. The entries of the associated confidence matrix were defined as a normalized distance of knockout levels to this threshold, see Supplementary Material S2. We then applied our response logic approach to these matrices to infer sparsified networks, as defined earlier. The goal of the DREAM challenge is to provide a list of gene pairs that is ranked by their predicted likelihood to be interacting. We generated it by first listing the predicted interacting and then the non-interacting gene pairs, where within each group, the pair list was ordered according to the associated entries in the global response matrix (interaction *i → j* was ranked higher than *k → l* if *R_ij_* > *R_kl_*). As comparison, we also created a ranked list by simply ranking gene pairs in the order of the global response matrix, without the grouping that was introduced by the response logic logic, which we termed “naive approach”.

These ranked lists were then scored using the official DREAMTools package [9] (with a minor modification for one network score at DREAM3 N=100, see Supplementary Material S2). Figure 4 shows the results of our method and that of the naive approach in comparison to the ten best performing participants at each network size and challenge. Except for the small networks with *N* = 10, where the response logic approach ranks second and third, it outperforms all 29 competitors participating in DREAM3 [31], as well all 29 competitors participating in DREAM4 [10]. Note that the best performers for the small networks (*N* = 10) that scored higher than the response logic [46, 28] also used the provided time course data, which we did not use in our response logic approach.

**Figure 4:**
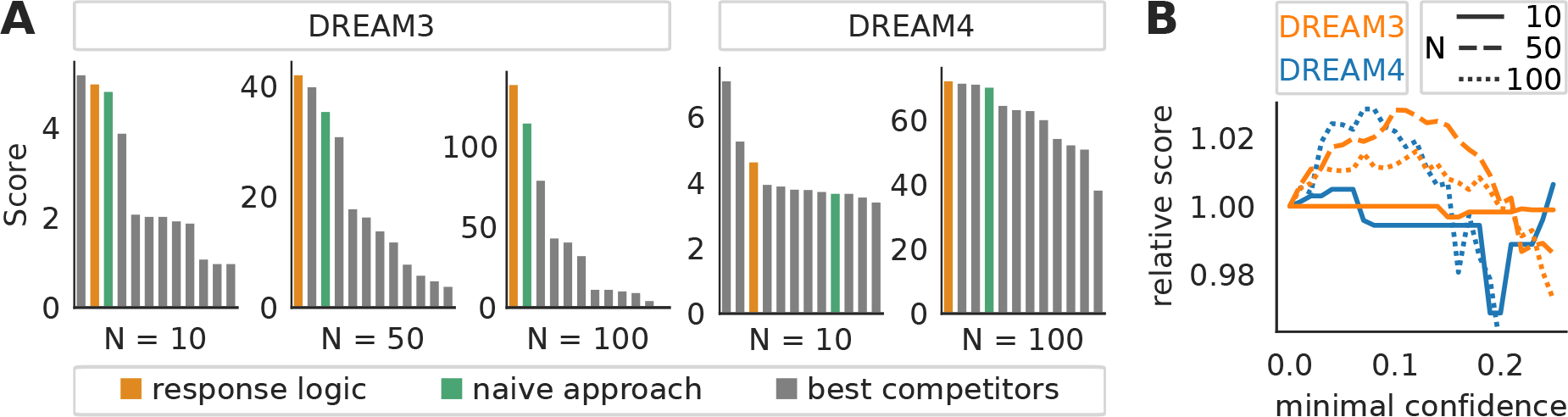
**A** Performance of the response logic approach for the gene-network reverse engineering challenges DREAM3 and DREAM4 [31, 20] (yellow bars), compared with a “naive” scoring approach (green) and the 10 best approaches that took part in the respective challenges. Scores are calculated as in the original challange, with higher scores indicating better performance. **B** Relative changes in performance when excluding data points with confidence below a certain threshold. *N*: network size

We also observed that the response logic always outperformed the naive approach, confirming that nontrivial additional knowledge is gained when applying the response logic. Notably however, already the naive approach scores comparatively well, which let the challenge’s organizers to conclude that “sophisticated methods that would in theory be expected to perform better than the naive approach described above, were more strongly affected by inaccurate prior assumptions in practice” [31]. This observation affirms our initial motivation to design an approach with minimal assumptions on the data.

Finally, the DREAM data also allowed us to test if disregarding low-confidence data points, as suggested by the KEGG pathway analysis, improves predictions. Thus, considering the confidence matrix with scaled entries between zero and one (Supplementary Material S2), we removed data points with confidence scores below a threshold and re-engineered the networks from those smaller data sets. The resulting scores relative to the original scores, which were obtained from the full response patterns, are shown in Figure 4B. With the exception of the *N* = 10 networks, these numerical experiments confirmed that removing low-confidence data effectively improved network inference. Peak performance is reached when approximately five percent of the data is discarded.

In summary, our benchmarks using the DREAM insilico challenges provide a strong indication that the response logic approach is capable of reverse-engineering biological networks. Its simplicity not only makes its results comprehensible but the DREAM challenge showed that they are also more accurate than those of existing methods.

### 3.3 Reverse engineering MAPK and PI3K signalling in a colon cancer cell line

After having benchmarked the response logic formalism, we next aimed to use it to investigate signalling networks in cancer cells. In a first step, we decided to reverse engineer the Ras-mediated signalling network including MAPK and PI3K/AKT signalling in SW-48 colon cancer cells. We performed multiple perturbation experiments using either ligands or inhibitors that targeted EGFR, PDFR, ERK and AKT, and measured changes in phosphorylation using a reverse phase protein assay (RPPA) platform. Ten of the antibody-based readouts passed a quality control and were relevant to the considered pathways, see details in Supplementary Material S3. Using replicate measurements of both unperturbed and perturbed conditions, we constructed the response pattern as well as the according confidence scores, which are shown in Figure 5A (see Supplementary Material S3 for details).

**Figure 5:**
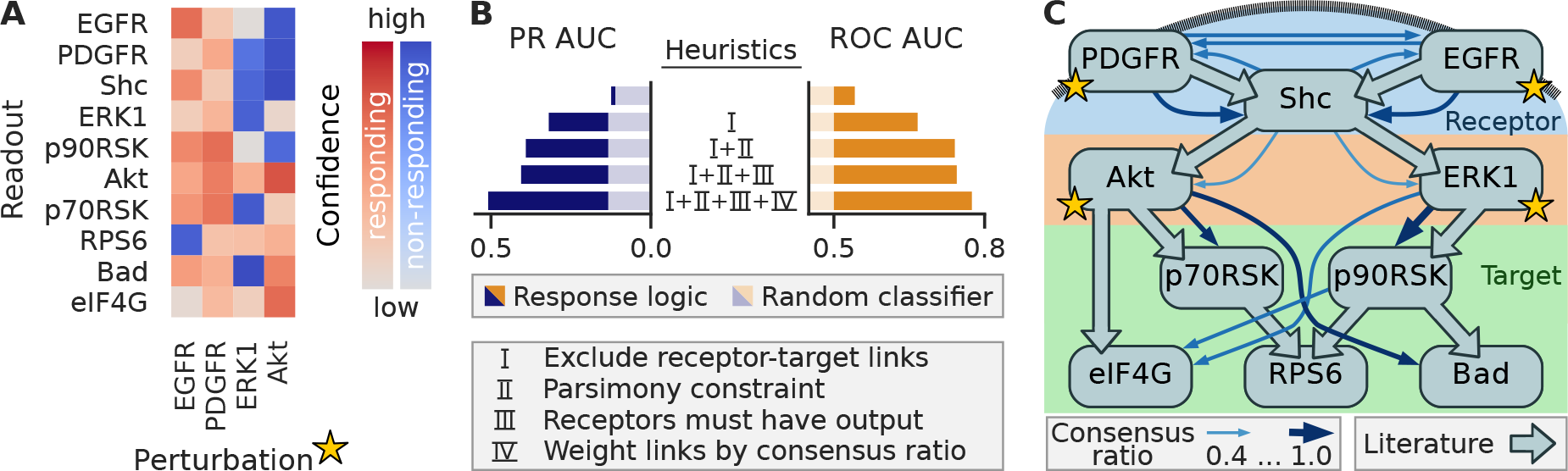
**A** Response pattern of the SW-48 cell line of selected phospho-proteins after perturbations affecting EGFR, PDGFR, ERK1 and AKT. **B** Performance of response logic network inference under various (combinations of) heuristics, as explained in text, compared to a random classifier (shaded colors). **C** Literature network (filled arrows) and final network prediction (finer arrows, only links with consensus ratio *≥* 0.4 are shown).

The RAS signalling network has been well studied, which allowed us to compile a literature network shown in Figure 5C that can be used as a gold standard to measure prediction performance. We then applied our response logic framework to the response pattern, and evaluated predictions by means of the areas under the ROC-, as well as precision-recall (PR) curves, as shown in Figure 5B, see Supplementary Material S3 for details. As it was computationally impossible to enumerate all networks, we determined the union of all conforming networks, as described earlier, and scored links based on whether they are found in all, in some and in no conforming networks. Doing so led to PR and ROC curves that were only marginally better than random (top row in Figure 5). The apparent challenge concerning the network inference for this network is the substantial disparity between ten readouts to only four perturbations, making the reverse engineering problem strongly underdetermined. A crucial benefit of the response logic analysis is that it allows allow for the incorporation of various additional insights about the structure of signalling networks to reduce the space of solutions. We therefore next investigated how the inclusion of generic and indirect network knowledge rendered the analysis more meaningful. First, we enforced a hierarchy in the network (heuristic I). Signalling networks typically process signals received on the receptor level through a chain of intermediate kinases, before they are passed on to a set of targets. We therefore disallowed any direct connections between the receptor and the target level (according to the allocation shown in Figure 5C) (these ruled-out links were obviously not taken into account for the scoring procedure, which explains the different areas under the precision-recall curve for the random classifier in Figure 5B). Furthermore, kinase interactions are highly specific, resulting in sparse signalling networks. Therefore, we found it reasonable to rid the network of redundant links and apply the parsimony constraint, as defined earlier (heuristic II). Lastly, we required that any node at receptor level must have at least one outgoing link (heuristic III).

Adding these three heuristics, I – III in Figure 5B, considerably improved the performance and reduced the solution space to 666 conforming networks. This makes it possible to enumerate them all and compute for each possible link the fraction of how many times it was present in all conforming networks (consensus ratio). We reasoned that a higher consensus ratio also implies a higher relevance, which we found confirmed when using the consensus ratio, rather than the union of networks to score the predictions (heuristic IV). From these results we conclude that the response logic is indeed a valid assumption for the MAPK and PI3K pathway activity in the SW-48 cell line and that rather apparent additional information can effectively compensate for the small number of perturbations.

### 3.4 Modelling the effects of PI3K mutations in a colon cancer cell line

Having verified the validity of the response logic approach on the SW-48 cell data, we next used it to investigate how different mutations in the PI3K change signalling. To investigate this, we used clones of SW-48, in which two mutations that are commonly found in tumours were integrated, namely PIK3CA^H1047R^ and PIK3CA^E545K^. We generated data using the same scheme as before, by perturbing the cells with ligands and inhibitors, and measuring the response using RPPA. Considering that the MAPK and PI3K pathways are very well studied, we assumed that the literature network depicted in Figure 5 is valid for all three cell lines, except for those links that could be affected by the different PI3K mutations. Because PI3K is not amongst the readouts, we model PI3K mutations to potentially affect links from and to its next downstream target, which is AKT. Furthermore, the literature network does not include context-dependent feedbacks in the MAPK pathway [29]. As we observed mutant-dependent upregulation of EGFR as well as SHC upon MEK inhibition, see Supplementary Material Figure S4, we considered this option in the inference as well.

Therefore, to model the different mutant response patterns Figure 6A, we used a heavily constrained response logic approach in which the presence or absence of network links is governed by the literature network, expect for those links going in and out of AKT and those links going into EGFR. Not only did these constraints compensate for the few perturbations but also connect differences in the data to plausible alterations of the network. Furthermore, as the parsimony constraint has proven beneficial in the response logic validation on the parental cell line data, Figure 5, it is used as well (with the constrained literature network, previous heuristics I and III no longer apply, and IV is not relevant as shown next).

**Figure 6:**
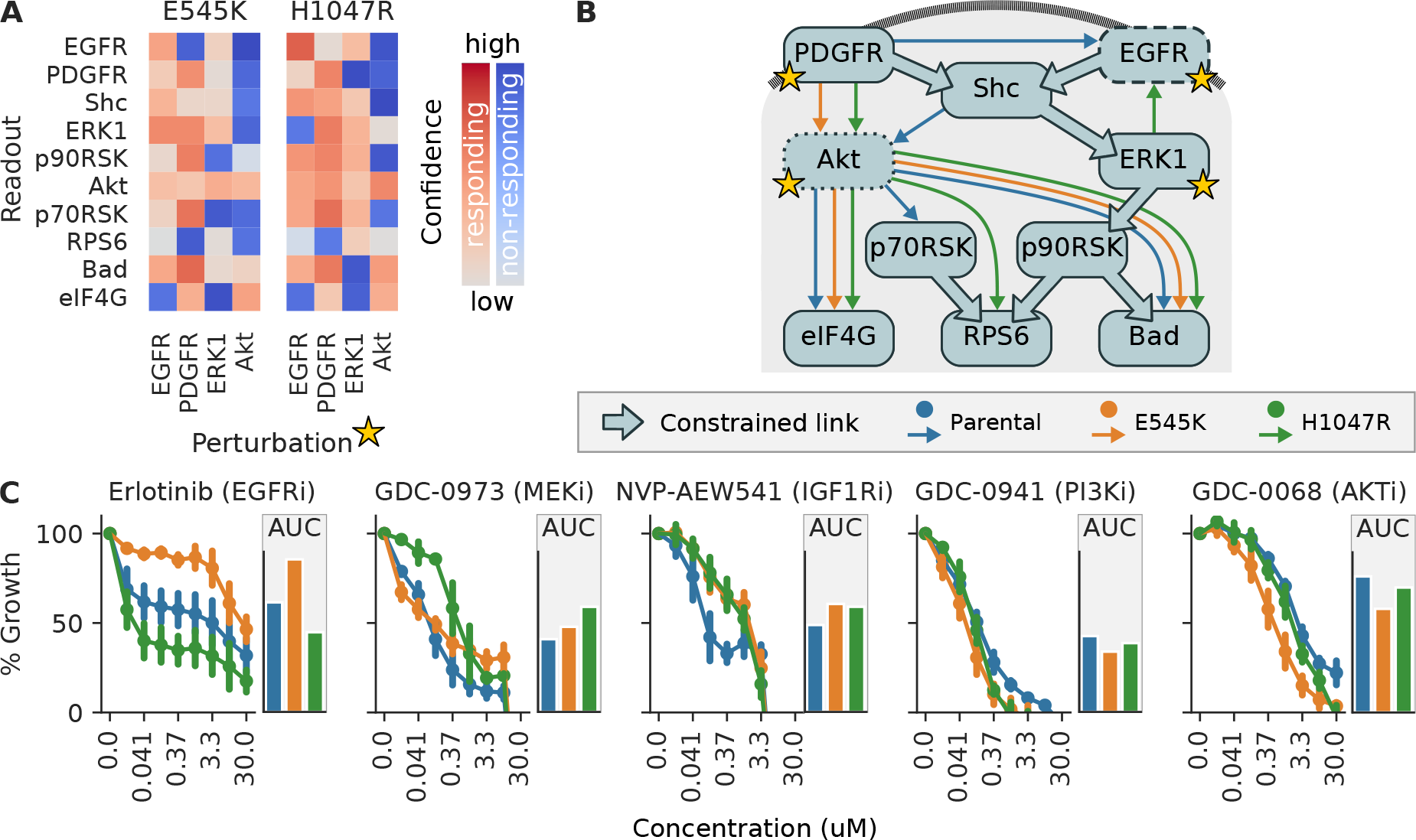
**A** Response pattern for two PI3K-mutant cell lines derived from SW-48, carrying either the E545K or H1047R mutations in PIK3CA, as in Fig. 5. **B** Mutant-specific networks derived from the response data (arrows in blue (parental cell line), yellow and green (two mutant cell lines), with links constrained due to literature knowledge shown with large arrows. **C** Dose-response curves for different inhibitors targeting the inferred networks in the parental cell line (blue) and the two clones with PI3K mutations (yellow and green), and the area under the curve displayed as bar graphs.

This approach resulted in four, one, and two conforming networks for the parental, the E545K, and the H1047R cell line, respectively. For the two ambiguous cell lines, we decided to isolate the single, biologically most plausible network hypothesis. In the case of the parental cell line, the four conforming networks consist of the combined options of whether or not SHC feeds back to EGFR and whether EGFR signals directly to AKT or via SHC. SHC has been found to be an adapter protein that is recruited to the activated EGFR (but does not activate it) and is essential for the receptor’s signal relay [35]. We thus chose the parental network hypothesis that excludes the SHC to EGFR and the EGFR to AKT link. The two H1047R networks only differed in whether a feedback to EGFR originates from p90RSK or from ERK. Since the ERK to EGFR feedback is well described in the literature [29], we decided for that option. With this, we could compare the mutant specific network hypotheses, as shown in Figure 6B, which led to two main observations. First, in contrast to the parental cell line, the mutant cell lines do not have a link from the EGFR receptor to the PI3K pathway. And second, the H1047R cell line is the only one bearing a feedback from ERK (or any node) to EGFR.

We next aimed to explore if these different network topologies might explain phenotypic differences between these cell lines. We therefore evaluated the drug-response of these cells for different targeted drugs, as shown in Figure 6C. Some differences in drug response can be understood directly from the mutations that have been added to the cell lines: The PI3K and AKT inhibitors seem to be slightly more effective in the PI3K mutant cell lines, which is not surprising as these cells have constitutively active PI3K signalling. Similarly, inhibition of IGFR was more effective in the wild type cells, as the mutant cells are more self-sufficient in PI3K signalling and therefore potentially require less IGFR activity. The drug responses to the EGFR inhibitor, and the MEK inhibitor are more complex and can only be interpreted when considering the network rewiring. The PI3K^H1047^ mutant cell line is rather re sistant to the MEK inhibitor. This can be understood by the presence of the negative feedback from ERK to EGFR in this cell line, which is known to cause resistance by re-activating ERK and amplifying AKT signalling upon MEK inhibition [27]. EGFR inhibition effects the cell line with the E545K mutant less, and the cell line with the H1047R mutant more strongly compared to the parental cell line. Both mutants decouple the EGF-receptor to the AKT pathway, so one would expect that they also show a less pronounced effect upon its inhibition. However, in the H1047R cell line there is a strong ERK-EGFR feedback that generally reduces the MAPK pathway activity, and one can speculate that additional EGFR inhibition can push the MAPK pathway activity to sub-critical levels.

Taken together, the response logic modelling allows to reconstruct networks from complex perturbation data and provides network information that can be interpreted and linked to phenotypic behaviour. This example demonstrates how this approach allows to integrate noisy response data, prior network knowledge and generic signalling constraints to identify hypothesis on changes in networks due to mutations, that can subsequently be studied experimentally.

## 4 Discussion

We developed the response logic approach as a method to infer directed networks from perturbation data. Its central idea is to assume that a perturbation of a node is propagated along the edges and thus causes a response at all nodes to which there is a directed path, starting from the perturbed node. This simple hypothesis is integrated in a logic program that allows to identify the networks whose transitive closure most closely matches that of the experimental data. The power of logic programming, and more generally declarative programming, has enabled its use in a wide range of topics in computational biology [14, 47, 42, 5, 6, 2]. In our approach, logic programming provides a way to efficiently scan the vast search space of all directed networks and to easily express and incorporate additional information and prior knowledge about the network.

Many reverse-engineering methods involve tunable parameters, which can drastically affect the results. However, it is often far from obvious how these parameters should be set in a specific context. In contrast, our response logic approach is parameter free and strictly infers the networks that follow from the provided response pattern and any additional constraints provided.

At first glance, it might seem wasteful to reduce the data to a binary information of responding versus nonresponding, when many experimental techniques allow to quantify the magnitude of response of the observed components. However, data binarization renders inference more general and robust, and in many settings, technical issues such as measurement error, heterogeneous data sources, or various normalization steps, make the interpretation of magnitudes difficult.

The idea to map an experimentally observable response pattern onto a transitive closure has been proposed before. It was hypothesised that the sparsest directed acyclic graph whose transitive closure matches the observed response pattern describes the direct regulatory interactions in gene networks [43]. Such a graph is also known as the transitive reduction and can be computed efficiently [1]. This approach was heuristicly expanded to also allow for some cycles, and to refine the inferred network by incorporating double mutant perturbations and information about up- and downregulation [41]. Yet, this procedure has several shortcomings: It cannot incorporate existing domain knowledge, it cannot handle missing data points, but simply considers an unknown or uncertain response behaviour as non-responding, and it only finds a single, most parsimonious, network, which might not necessarily represent the underlying structure.

This last point is a strategy to compensate for the fact that network inference is an inherently underdetermined problem, because the number of independent measurements generally falls short on the number of possible interactions [12]. The response logic approach explicitly addresses this problem as it considers the entire ensemble of conforming networks rather than to single out a particular one, based on some fixed and potentially inaccurate assumption. It thereby reveals which parts of the network can not be inferred from the information provided so far. This important insight can then be used to either guide additional experiments or to systematically reduce the solution space by adding constraints that are most warranted in the given context. We consider this a crucial advantage over existing approaches, whose inferred networks can generally not be intuitively traced back to the data and thus tend to disguise if and how the inferred network is justified by the data.

But while the response logic is based on a simple and intuitive concept, such simplicity comes at a price. As with any other assumption, it might not actually be representative of the studied system. Major problems might occur due to robustness, or saturation effects, all of which disrupt the presumed flow of signal but are an essential part of various biological systems [15]. Also, from a boolean perspective the response logic treats nodes exclusively as OR gates, whereas certain systems require a more involved logic [36]. Another important shortcoming for many questions is that it does not assign any weights nor signs to the inferred links. Yet, the inferred network can serve as an input for methods that are devised to quantify link strengths on a given input network [13].

On the other hand, the response logic’s simplicity makes it suitable for various different fields of research. Because it is based on a formalization of an intuitive network behaviour, it can infer ecological, infection, or even social or transportation networks. Such generality would even permit to use the response logic to ask the inverse question: Given a certain network structure and the observed perturbation responses, can I infer where a perturbation hit the network? This question could be particularly interesting in the analysis of man-made networks, for which the structure is typically known, but not the location of the perturbation. The inverted logic program would then identify where an electric connection malfunctioned, an intruder attacked, or a disease originated from. All these possibilities show that the simplicity of the response logic does not limit its applicability.

## Acknowledgements

The authors would like to thank Dr. Manuela Benary, Mathurin Dorel, Dr. Bertram Klinger and Dr. Johannes Meisig for fruitful discussions.

## Funding

This work was supported by a grant from the Berlin Institute of Health and by the German Research Foundation (DFG), through GRK1772.

## S1 Supplement

### S1.1 Encoding the response logic in ASP

The response logic is formulated in the framework of Answer Set Programming. We use the syntax of the input language gringo. A straightforward introduction is provided in section 4 of [42]. The following facts and clauses constitute the logic program.

At the beginning, the program needs to state the facts, i.e. the number of nodes in the network as constant n, the set of perturbations as functions `pert(I,OUT)`, where variable `I` names a specific perturbation and variable `OUT` indicates the node(s) targeted by the perturbation (could be multiple), as well as the rectified response pattern as functions `response(I,N)` and `-response(I,N)` (classical negation), where variable `I` names a specific perturbation and variable `N` indicates the node(s) responding or not responding to the perturbation, respectively.

The program then follows the generate-define-test pattern, as below. The function `tc(I,IN)` represents the transitive closure of the answer set network.

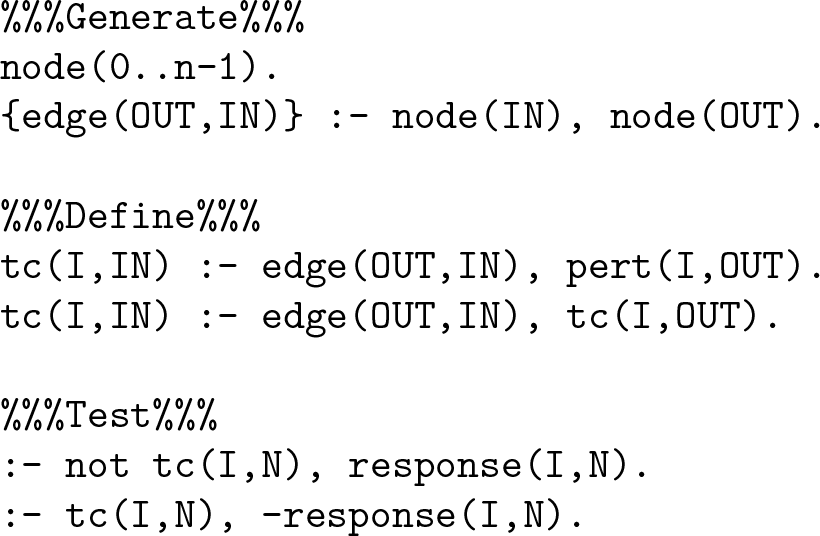

The last two lines are the integrity constraints that discard any answer set whose transitive closure is in disagreement with the rectified response pattern. ASPs distinction between classical (”-”) and default (”not”) negation allows to only test the integrity for the known entries of the response pattern and to ignore the test for any entries that are not known.

For large networks the search space of the above program becomes very large with negative effects on performance. This is especially problematic during the rectification of the response pattern because in principle every entry of the response pattern must be tested for consistency. We implemented two strategies to speed up the process. First, at each step of the iteration, we check if the previous transitive closure is in agreement with the new data point. If so, we know that there is a conforming network and can accept the data point without an explicit test. Second, if there is no additional constraint (see below) stating the absence of a network edge, we can significantly simplify the logic program. In that case, if the logic program is satisfiable at all, the network with directly links from perturbed node to all its responding nodes, i.e. a network with star topology, constitutes a conforming network. As the rectification process only needs to determine satisfiability, a much more simple logic program avoids generating all possible networks but only tests the star network (which we do not need to state explicitly):

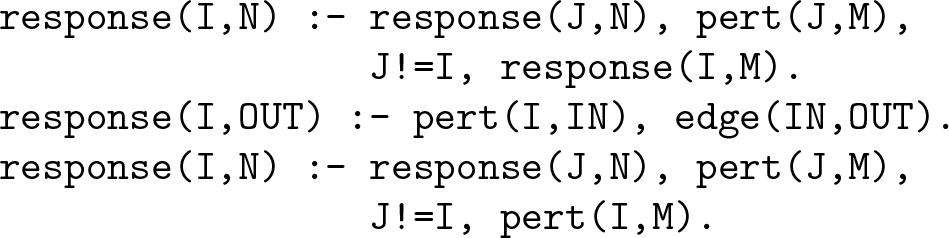

The idea of this encoding is to define all additional responses that are implied by the star network from the given response pattern (line 1): if node N responds to a perturbation of node M (it is reachable from M) then node N will also respond to any other perturbation for which it is known that it causes a response at node M. This defines the minimal set of responding nodes consistent with the response logic. Any constraints on the absence of edges would imply (longer) non-direct paths that could potentially imply more nodes to respond to certain perturbations. This program does not even need to explicitly state an integrity constraint because an observed non-response given as classical negation `-response(I,N)`, already rules out that `response(I,N)` is contained in the (single) answer set. Furthermore, line 2 adds responses that are implied by edges that are known to exist and the last line is needed in the case where multiple perturbations hit the same node, but a perturbed node itself does not respond.

As discussed at length, we might wish to incorporate additional, external domain knowledge into the logic program. The most direct way of doing so is to simply state the knowledge about presence or absence of e.g. the link from node 2 to node 3 as a fact:

~~~
edge(2,3).
~~~

or

~~~
−edge(2,3).
~~~

respectively.

To encode bounds on the number of links that enter or leave one or a group of nodes is simple, due to ASPs built-in `#count` aggregates. If, for example, we were to maximally allow for 3 incoming edges to node 1 we would add the following integrity constraint to the logic program

~~~
:− #count {OUT : edge(OUT,1) } > 3.
~~~

Obviously, this statement could easily be adapted to formulate a lower bound, or bounds for groups of nodes.

It is more complex to state the parsimony constraint that filters out any network for which the removal of any link does not change the transitive closure. Essentially, for a given network, we define a set of links, x edge(OUT,IN) that are subject to removal. Those are all the edges that are part of the network but which are not enforced by external knowledge. Then, the idea is to define all networks with a link removed, determine their transitive closure and compare it to that of the original network. The final integrity constraint removes the answer set if any of the reduced networks does not have a reduced (that is, it has the same) transitive closure as the original one.

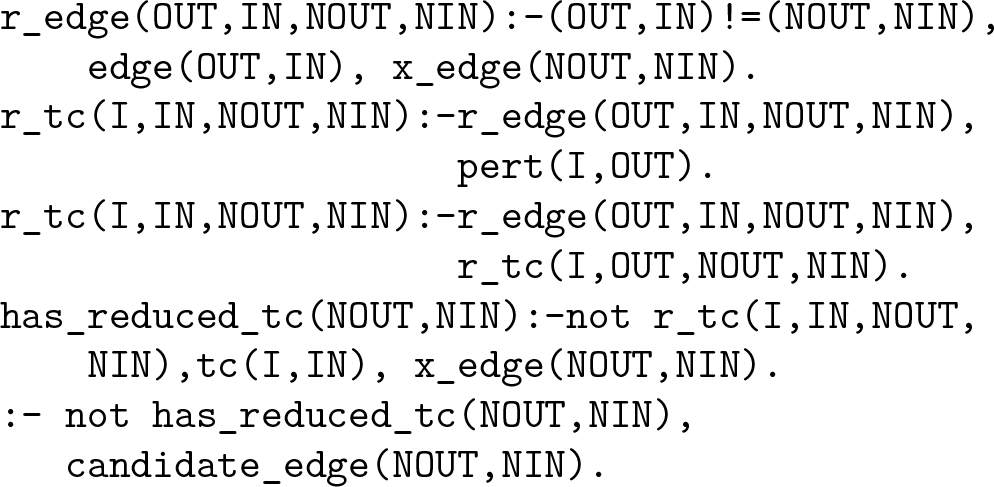

As this logic creates a significant amount of additional variables, we observed that the parsimony constraint will suffer performance issues when applied to more than a few tens of nodes.

Finally, a crucial feature of the response logic approach is the ability to generate the union of all conforming networks, which points out the links than can or cannot be uniquely determined. Typically, the number of conforming networks is intractable which precludes to simply enumerate all solutions and then compute their union directly. Rather, we let a logic program find it directly. The union of all answer sets of an ASP program is also termed brave consequences, hence the naming conventions below. In a first step, we need to annotate the presence or absence of an edge explicitly.

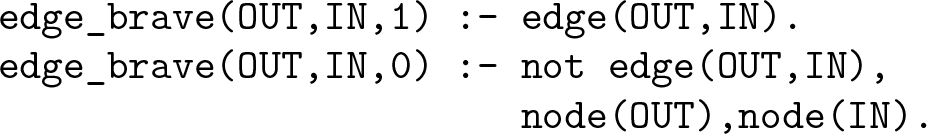

Then, the idea is to repeatedly solve the logic program, while iteratively building up the union of conforming networks. That is, we record for each edge whether the solutions generated so far, found it to be always absent, always present or neither (sometimes present, sometimes absent). To obtain the union over all answer sets, the program is further constrained with every new solve call to only permit networks that are not a subset of the union that was recorded so far. This is encoded by the following integrity constraint.

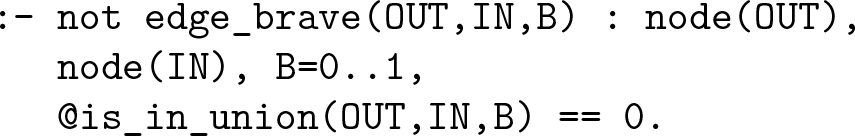

The @is in union(OUT,IN,B) construct is a function called by the solver that will return 1 if the presence or absence (`B` is 1 or 0, respectively) of edge `OUT` to `IN` is already recorded in the union and 0 otherwise. Thus, every solve call adds new elements to the union until there are no more conforming networks that show the presence or absence of a link, beyond what is already represented in the union. This is when the program is no longer satisfiable and the repeated solve calls stop.

### S1.2 Additional information about the inference of DREAM in-silico networks

The rectification of the response pattern requires confidence scores, *C*. Those are defined as a normalized distance of each entry of the global response matrix *R* to 1, which was chosen as the response threshold. The normalization is chosen such that confidence levels range from zero, for *R_ij_* = 1, to one, for *R_ij_* taking either the maximal *R*^max^ or minimal *R*^min^ value of all entries. Formally,

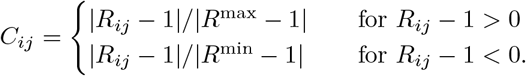

Both DREAM3 and DREAM4 provided five network challenges per network size. The precision-recall curve and the ROC curve for each network prediction are shown in Figure S1 and Figure S2.

The official DREAMTools package [9] was used to compute the final prediction scores that are shown in main text Figure 4A. The scoring is based on p-values for the ROC and precision-recall area under the curves (AUC) that are computed from probability distributions of random network predictions. In the case of the DREAM3, N=100, Yeast3 network the precision-recall AUC of the response logic prediction is larger than that of any of the provided random network predictions. Therefore the determined p-value becomes zero and the overall network score becomes infinity. To obtain a more reasonable score we decided to modify the p-value computation in that case. We identified the largest AUC for which the random probability distribution shows a nonzero probability and simply defined an extended distribution whose probability decreases linearly from this point to zero probability at AUC=1 and computed the p-value based on this approximated distribution. This only concerned one of the five scored networks in the DREAM3, N=100 prediction and the resulting overall score, 139.7, is shown in main text Figure 4A, third panel. Another strategy to cope with the problem is to remove the network altogether and compute the overall score (mean of geometric mean of pvalues) only from the remaining four networks. This provides a lower bound for the overall score of 122.9, which is still clearly better than the score of any of the competitors and the naive approach.

### S1.3 Additional information about the analysis of the SW-48 cell line

A reverse phase protein array (RPPA) perturbation experiment was carried out on the SW-48 parental, the SW-48 E545K and SW-48 H1047R PI3K mutant cell lines under full serum conditions. Antibodies were chosen to cover the activity (phosphorylation) of a range of kinases. Cells were perturbed by single small molecule inhibitors, by single growth factors or by a combination of one growth factor and one inhibitor. Measurements were carried out in 8 technical replicates.

In a first step readouts were filtered as follows. First, we removed all readouts whose signal remained within the technical noise level throughout the treatments, as this indicates a malfunctioning antibody. The readout had to be a component or a target of the MAPK or the PI3K pathway. To decrease redundancy, we removed very closely related readouts. This included readouts of functionally related phosphosites on the same kinase, or kinases that showed near-identical qualitative behaviour due to their proximity within the signalling pathway. This left us with 10 different readouts. Concerning the perturbations, we filtered out ineffective, or redundant inhibitors and ligands. This resulted in the data set depicted in Figure S3.

To use this data in the response logic framework we need to convert it to a response pattern with according confidence scores. First, we need to localize the targets of the perturbations. As not all the direct targets of each perturbation were part of the readouts, we replaced those by the ones that were the closest downstream the signalling chain. Concerning HGF stimulation, we observed a strong response by the PDGF receptor. C-Met not being amongst the readouts, we chose the target accordingly. Lacking additional data, it is subject to speculation whether this behaviour could be explained e.g. by receptor co-activation [45] or impurities of the used ligand. This resulted in the following allocations.

| Perturbation | Target |
| --- | --- |
| EGFRi | EGFR |
| MEKi | ERK |
| AKTi | AKT |
| EGF | EGFR |
| HGF | PDGFRB |
| IGF | AKT |

This makes apparent that the perturbations only have four different targets (PDGFRB, EGFR, ERK, AKT).

Next, we need to determine the response behaviour with respect to perturbations of those four targets. Inhibiting an unstimulated signalling pathway can lead to saturation effects, when additional reduction of kinase activity is not possible. Thus, to faithfully track the inhibition response we decided to investigate inhibition while cells are stimulated, that is to compare ligand + inhibitor to only ligand. To ensure that such a stimulation actually affects the inhibited pathways, we chose EGF stimulation for the EGFR and MEK inhibitors and IGF stimulation for the AKTi inhibitor. The stimulation effects do not suffer from such saturation effects and were thus compared to basal levels (DMSO+PBS) directly. We compiled an overview of the resulting perturbation comparisons in Figure S4.

To decide whether a perturbation caused a response, we computed p values for each pair of perturbed-unperturbed samples using an unpaired, two-sample t-test. As the data replicates are technical in nature we chose a conservative significance threshold of 0.01. To compute confidence scores we used the same procedure as for the DREAM data, subsection S1.2, that is, the confidence scores represent a normalized distance between the p-value and the p-value threshold. Recall, that the six perturbations only have four different targets. To remove redundancy in the response logic sense, we decided to remove the two redundant data points with lower confidence (per cell line and readout). This resulted in the response patterns shown in main text Figure 5A and 6A.

The computation of the area under ROC and precision recall-curves in main text Figure 5A requires to rank the predicted links. However, depending on how the response logic approach is applied, some links can end up with the same score. We therefore, computed the PR AUC and ROC AUC as a mean over 100 AUC values that were generated by randomly reshuffling groups of links with equal score. Repeated computation of these PR AUC and ROC AUC values only varied in the third decimal place.

**Figure S1:**
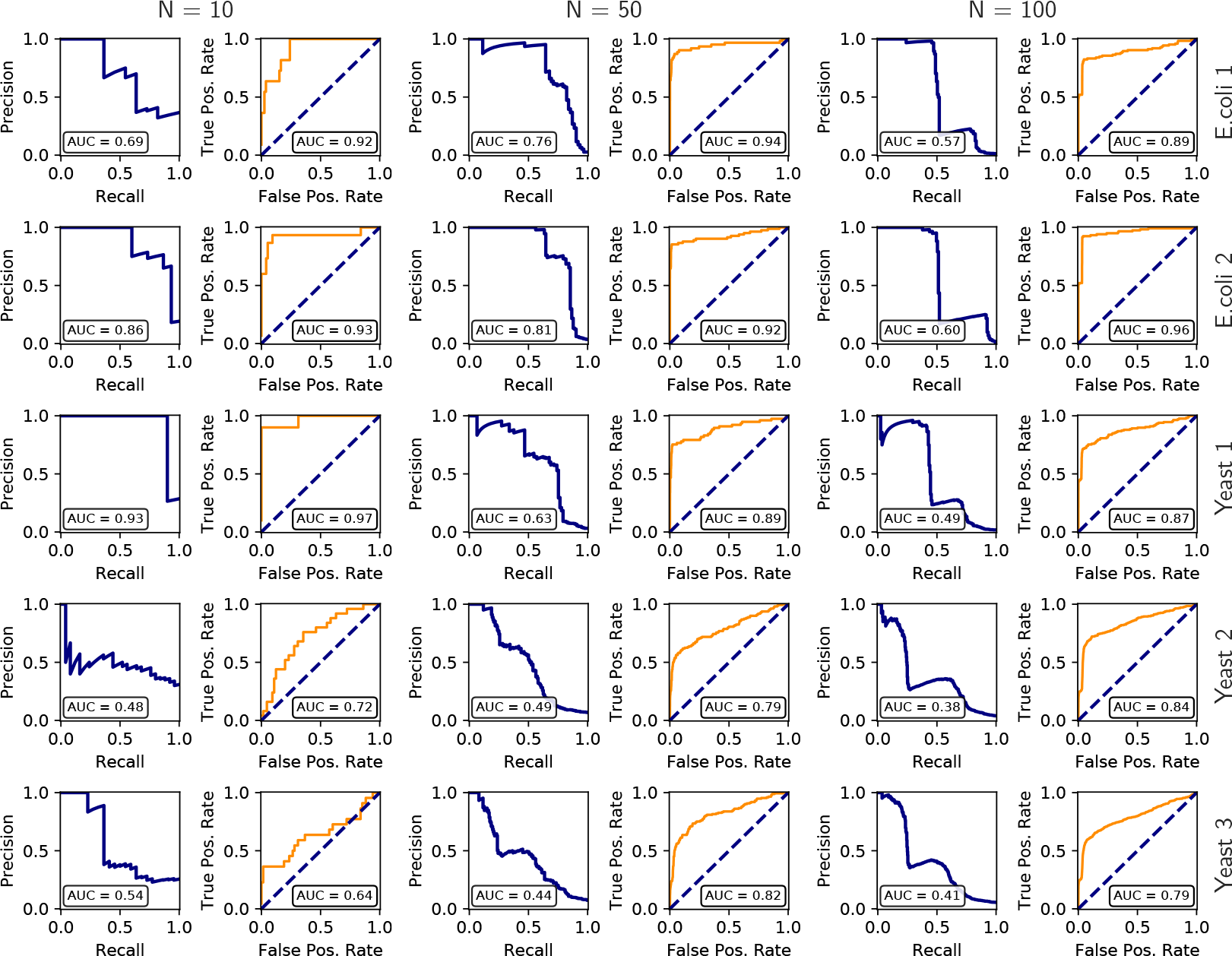
DREAM3 challenge 4: Precision Recall (blue) and ROC (red) curves for the response logic predictions of the five networks.

**Figure S2:**
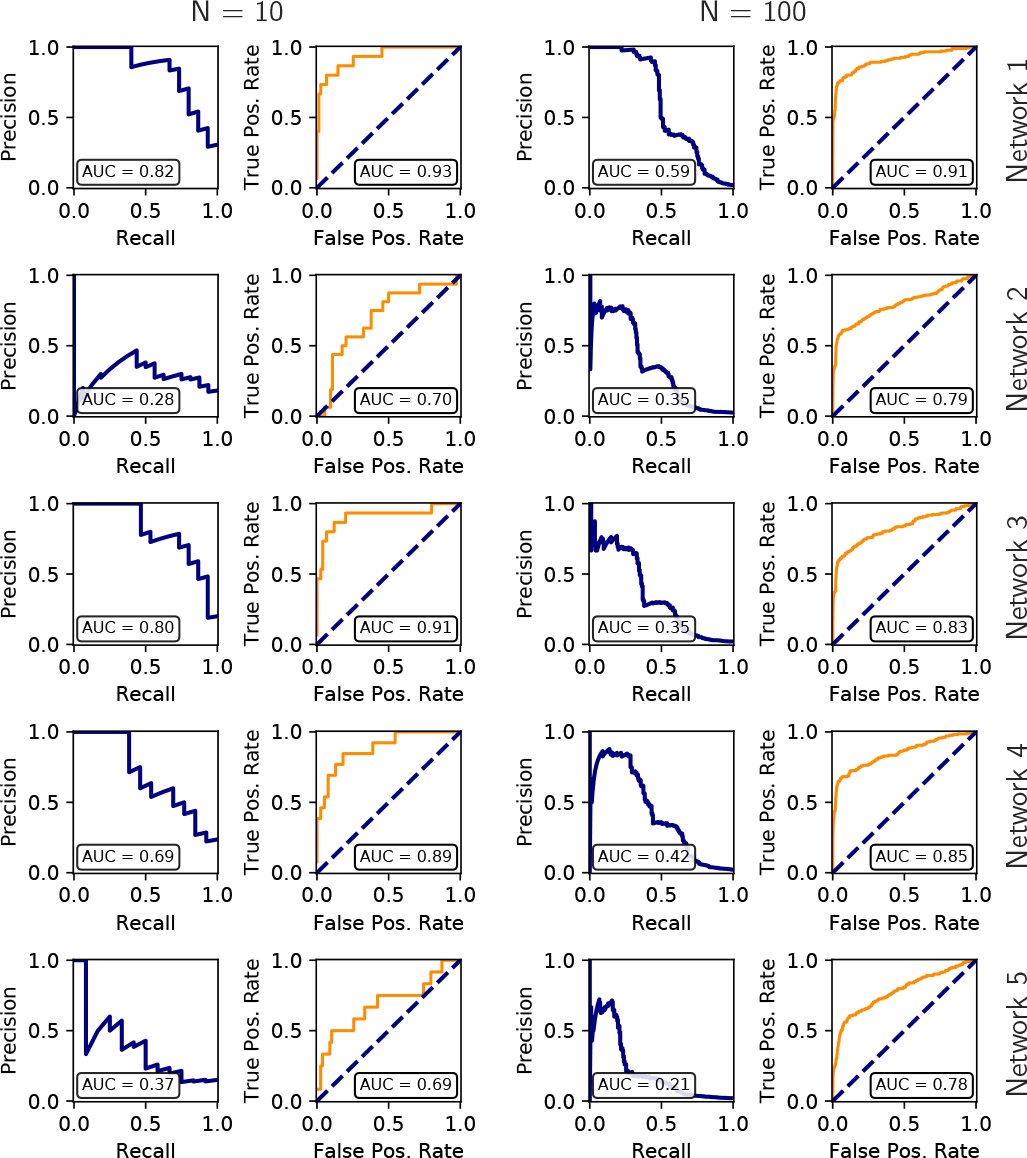
DREAM4 challenge 2: Precision Recall (blue) and ROC (red) curves for the response logic predictions of the five networks.

**Figure S3:**
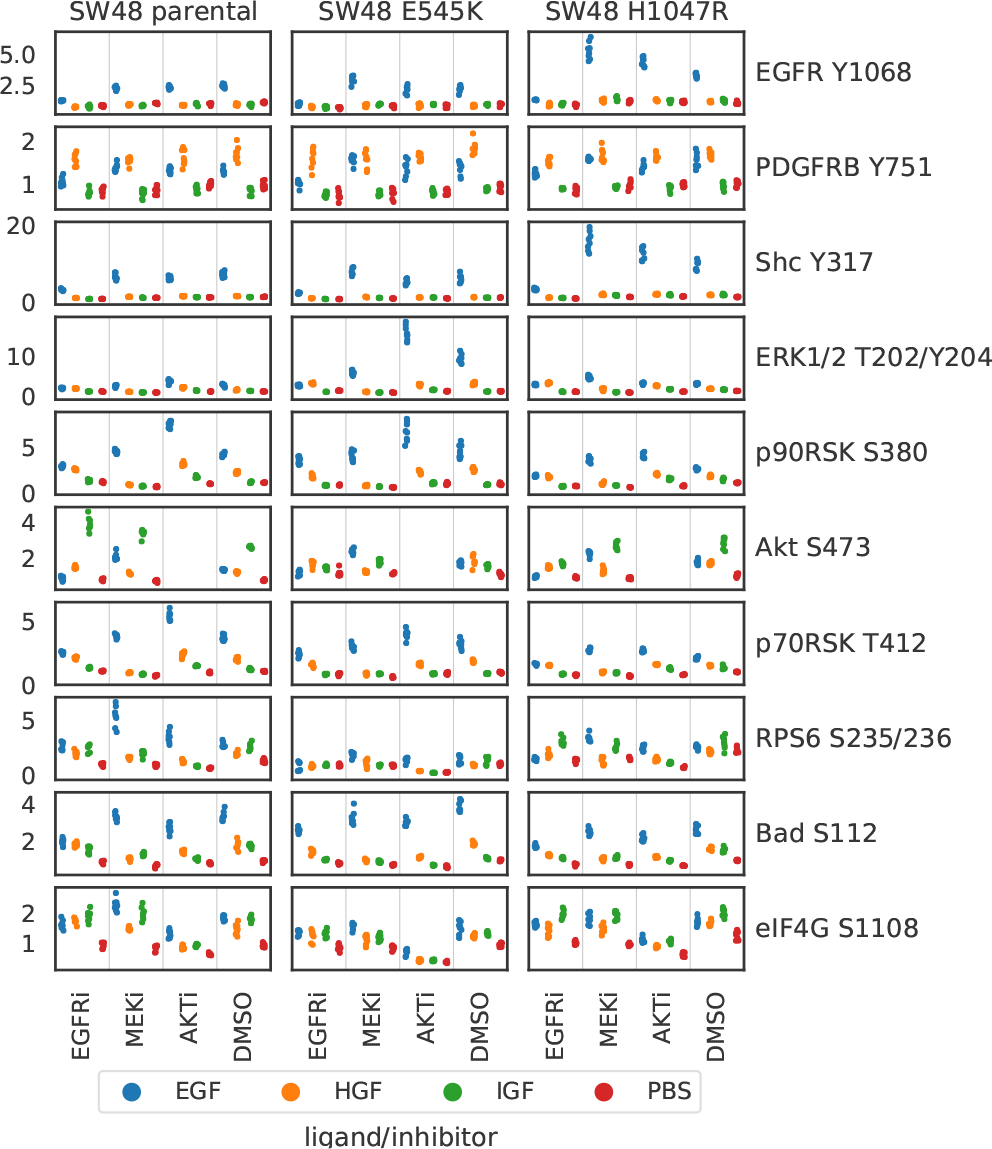
RPPA measurements on SW48 cell lines

**Figure S4:**
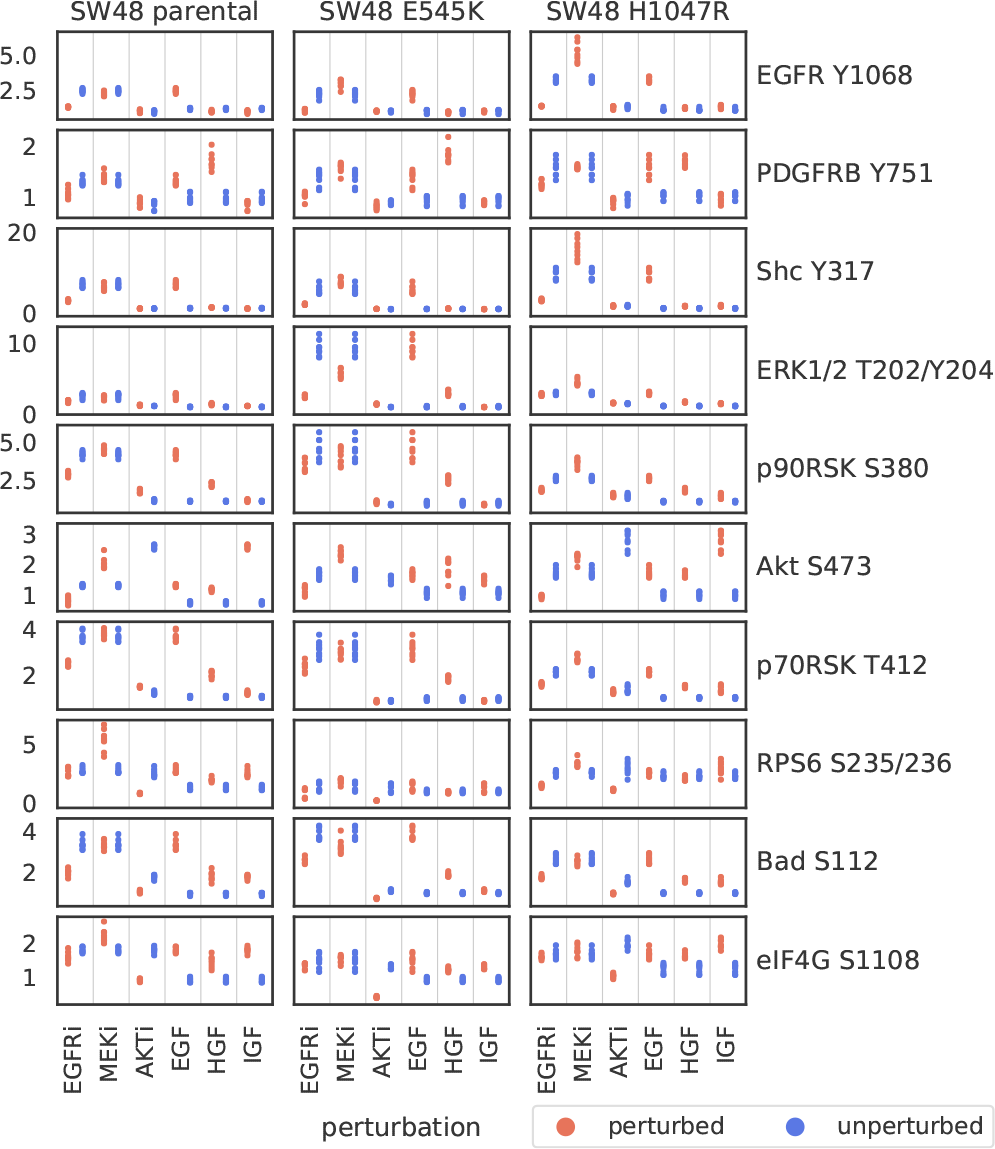
Same data as in Figure S3, but filtered and regrouped to highlight perturbation effects.

